# Modulation of prefrontal functional connectivity by anodal tDCS over left DLPFC predicts performance enhancement in competitive swimmers: A simultaneous tDCS-fNIRS, double-blind, sham-controlled crossover study

**DOI:** 10.1101/2025.07.11.664462

**Authors:** Toru Ishihara, Shoki Kiyokawa, Haruki Tajimi, Shinnosuke Hashimoto, Tongfang Ding, Kazuki Hyodo, Haruto Takagishi, Tetsuya Matsuda

## Abstract

While transcranial direct current stimulation (tDCS) has been proposed as a method to enhance physical performance in athletes, the underlying neural mechanisms and the reasons for the widely reported individual variability in its effects remain unclear. This study investigated whether prefrontal hemodynamic responses, measured by functional near-infrared spectroscopy (fNIRS), are associated with the effects of anodal tDCS over the left dorsolateral prefrontal cortex (DLPFC) on swimming performance. In a double-blind, sham-controlled, crossover design, eight trained male swimmers performed 100 m freestyle trials under both anodal tDCS and sham conditions. We recorded prefrontal cortical activation and functional connectivity using fNIRS during a resting-state period and a subsequent stimulation period. While tDCS led to a numerical improvement in 100 m freestyle time, the overall effect was not statistically significant. The fNIRS analyses revealed that tDCS significantly reduced intra-hemispheric functional connectivity, especially in the stimulated left prefrontal cortex. Crucially, the magnitude of this connectivity reduction correlated with the degree of performance improvement, suggesting a direct brain-behavior link. Exploratory analyses further suggested that baseline functional connectivity could predict an individual’s neural response to tDCS, with those having higher baseline connectivity showing a greater reduction. These findings suggest that tDCS over the left DLPFC may enhance physical performance by increasing the neural efficiency of prefrontal networks. Therefore, baseline functional connectivity is a promising physiological biomarker that could be used to personalize neuromodulation protocols in athletes.

## INTRODUCTION

Transcranial direct current stimulation (tDCS) is a non-invasive neuromodulation technique that alters cortical excitability and can modulate human behavior (1). It has recently gained attention as a potential tool for enhancing physical performance (2–4). However, findings remain inconsistent, with some studies reporting improved performance while others find negligible or no effects. Although recent meta-analyses suggest that a single session of tDCS may yield small-to-moderate performance enhancements (3, 4), an umbrella review concludes that current evidence is insufficient to support its efficacy as a reliable intervention (5). This uncertainty presents a major hurdle for the practical application of tDCS in sports, where reliable and predictable gains are paramount.

A likely source of this inconsistency is the substantial inter- and intra-individual variability in response to tDCS (6–11). For instance, only about 50% of individuals show the expected increase in motor-evoked potentials (MEPs) following motor cortex stimulation (6–8). Furthermore, intra-individual reproducibility tends to be low, with intraclass correlation coefficients often below 0.3 (9–11). These findings suggest that a measurable neural response may be a prerequisite for achieving performance benefits, highlighting the need to understand the factors driving individual differences in tDCS efficacy.

To date, however, few studies have attempted to link these individual neural responses directly to athletic performance outcomes. Functional near-infrared spectroscopy (fNIRS) is a portable, non-invasive neuroimaging method that allows for the real-time assessment of cortical hemodynamics. Compared to other modalities like fMRI or EEG, fNIRS offers greater ecological validity for studying brain-behavior relationships in more naturalistic environments, such as athletic settings. A systematic review identified only 28 studies combining tDCS and fNIRS, none of which focused on physical performance as the primary outcome (12). While existing fNIRS studies suggest that tDCS can modulate cortical activation (12), its effects on functional connectivity are debated, with reports of both increases (13–19) and decreases (17, 19).

This study investigates the neurobehavioral effects of tDCS in competitive swimmers—an elite population in which even marginal gains can have significant implications. We focused on the 100 m freestyle sprint, an event where outcomes are often decided by fractions of a second, making it a sensitive task for detecting subtle neuromodulatory effects. Although previous studies investigating the effects of anodal tDCS on swimming performance have focused on various loci (left temporal cortex, left dorsolateral prefrontal cortex [DLPFC], left orbital prefrontal cortex) as stimulation targets and yielded mixed results (20–22), prior studies have shown that anodal stimulation of the left DLPFC can improve or maintain performance in competitive swimmers (22). Further, concurrent tDCS–fNIRS studies have demonstrated that this same montage reliably elicits detectable changes in prefrontal hemodynamics (23). Therefore, we focused on the left DLPFC as the target locus.

To address this critical gap in the literature, the present study used a simultaneous tDCS-fNIRS approach to investigate the physiological mechanisms underlying performance changes. We addressed three key questions: (1) Does anodal tDCS over the left DLPFC enhance 100 m freestyle swimming performance? (2) How does tDCS over the left DLPFC modulate PFC activation and functional connectivity? (3) Is there a relationship between these neural changes and any changes in physical performance? We hypothesized that individuals exhibiting greater tDCS-induced changes in PFC activation and functional connectivity would also demonstrate more substantial improvements in swimming performance.

## MATERIALS & METHODS

### Participants

Nine male university swimmers from a competitive swimming team were initially recruited via convenience sampling. One participant withdrew due to an injury unrelated to the study, resulting in a final sample of eight swimmers. Eligibility criteria included no history of orthopedic, cardiovascular, endocrine-metabolic, neurological, or psychiatric disorders, in accordance with the Japanese safety guidelines for transcranial electrical stimulation (Japanese Society of Clinical Neurophysiology, 2019). All participants completed a screening questionnaire to confirm their eligibility, which included questions about prior episodes of fainting or seizures. They also provided demographic data, competitive history, training frequency, and their personal best time for a 100 m freestyle in a short-course pool. Participants were instructed to abstain from alcohol for 24 hours before each session and to avoid caffeine and intense physical activity on the day of testing. They were also asked to ensure they had sufficient sleep the night before. Written informed consent was obtained from all participants prior to the study. The study was approved by the Human Ethics Committee of the Graduate School of Human Development and Environment at Kobe University.

### Experimental Procedure

The experimental protocol is illustrated in Figure 1. Each participant completed two sessions—one with active tDCS and one with sham stimulation—separated by one week. The order of the conditions was counterbalanced across participants using a computer-generated randomization list. Both participants and experimenters were blind to the assigned condition (double-blind design). Upon arrival at each session, participants completed informed consent procedures, underwent body measurements (height and weight), and filled out pre-experiment questionnaires. The fNIRS probes were attached to the forehead, after which the tDCS electrodes were positioned and secured with elastic bands. After ensuring the participant was comfortable, a continuous 28-minute fNIRS recording began, consisting of an 8-minute resting-state baseline followed immediately by a 20-minute stimulation period (tDCS or sham). During the recording, participants were instructed to remain still and fixate their gaze on a cross (0.8 × 0.8 cm) displayed on a white screen placed 50 cm away. Following the stimulation, they completed a sleepiness rating scale, performed a self-paced 5-minute warm-up in the pool, and then executed a 100-meter freestyle time trial from a water start in a short-course pool. Lap times were recorded every 25 meters. The session concluded with a post-exercise cooldown and a safety and sensory perception questionnaire to assess blinding effectiveness and sensory perceptions.

**Figure 1.**
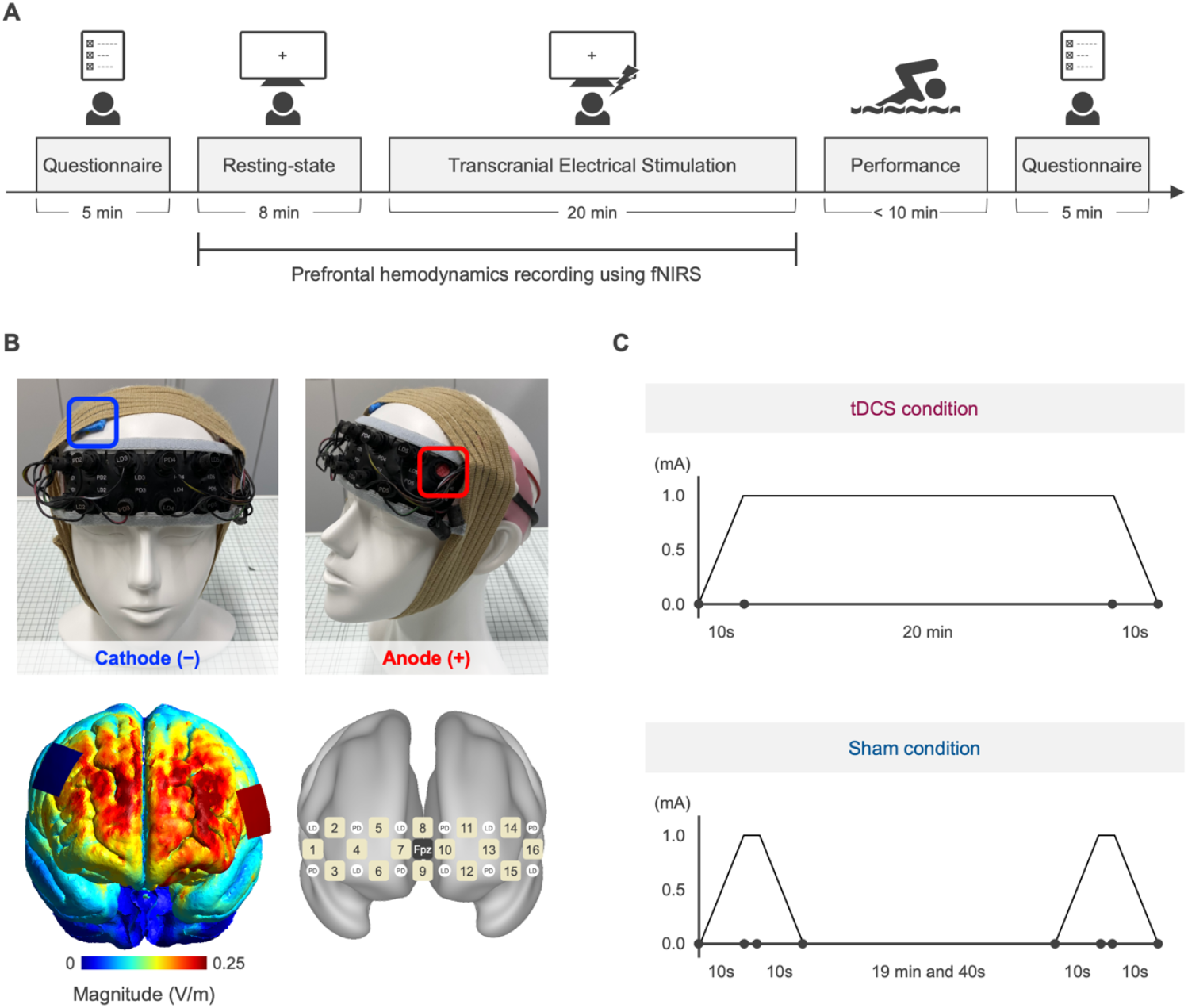
Experimental procedure and stimulation setup. A. Schematic of the experimental timeline. Participants first completed a pre-experiment questionnaire (5 min), followed by an 8-minute resting-state session with fNIRS measurement of prefrontal hemodynamics. This was followed by 20 minutes of either transcranial direct current stimulation (tDCS) or sham stimulation. Immediately after stimulation, participants performed a swimming task (100 m freestyle, <10 min) after 5 min warm-up, and finally completed a post-experiment questionnaire (5 min). B. Montage of the tDCS setup and current distribution simulation. The anode (+) was placed over the left prefrontal cortex (F5), and the cathode (−) over the right prefrontal cortex (F4), using 3 × 3 cm sponge electrodes. A headband was wrapped over the fNIRS optodes to shield them from ambient light during measurement. The lower left panel shows the simulated distribution of current density across the cortex. The lower right panel shows fNIRS probe placement over the prefrontal cortex. C. Stimulation protocols for tDCS and sham conditions. The tDCS condition applied a constant 1.0 mA current for 20 minutes. In the sham condition, current was ramped up and down briefly at the beginning and end to mimic the sensation of stimulation, but no sustained current was delivered.

### Transcranial Direct Current Stimulation (tDCS)

tDCS was delivered using the DC-STIMULATOR Plus (NeuroConn, Germany). Two 3 × 3 cm rubber electrodes were used, enclosed in saline-soaked sponges. The anode was positioned over the left prefrontal cortex (F5), and the cathode over the right prefrontal cortex (F4), based on the international 10–20 EEG systegm. Electrode placement was informed by simulations in SimNIBS 4.0 (24) to optimize current flow through the fNIRS measurement regions at 1 mA stimulation (Figure 1B). In the active tDCS condition, the current was ramped up to 1 mA over 10 seconds, maintained for 20 minutes, and then ramped down over 10 seconds (Figure 1C). In the sham condition, the same ramp-up and ramp-down sequences were applied at the beginning and end of the 20-minute period, but no current was delivered in between (Figure 1C). This procedure is a standard method for maintaining participant blinding.

### fNIRS Data Acquisition

Prefrontal cortical hemodynamics were recorded using a 16-channel fNIRS system (Spectratech OEG-16; Spectratech Inc., Tokyo, Japan) with two wavelengths of near-infrared light (770 and 840 nm) at a sampling rate of 0.76 Hz. The probe set was positioned symmetrically on the forehead, with its center aligned to Fpz (10–20 system). The source-detector separation distance was 3 cm for all channels. Participants were instructed to minimize head and facial movements during recording. Channels 14 and 16, which were located directly under the tDCS electrodes (Figure 1B), were excluded from the analysis to avoid potential artifacts.

### fNIRS Data Processing and Analysis

fNIRS signals were processed using proprietary software (OEG-16 v3.0) and R (version 4.5.0). Concentration changes in oxyhemoglobin (oxy-Hb) and deoxyhemoglobin (deoxy-Hb) were calculated using the modified Beer–Lambert law (25). A hemodynamic separation method was applied to reduce systemic physiological noise, such as skin blood flow, and to more effectively isolate neuronally-derived signals (26). Subsequent analyses focused on the oxy-Hb signal, which typically has a higher signal-to-noise ratio and greater sensitivity to cortical activation (27). A low-pass filter with a cutoff frequency of 0.1 Hz was applied to the oxy-Hb time series to attenuate physiological artifacts such as cardiac pulsation, respiration, and Mayer waves. The filtered data were then z-score normalized using the mean and standard deviation of the 8-minute resting period for each channel. Given the slow nature of tDCS-induced hemodynamic responses, baseline drift correction carries the risk of inadvertently removing or distorting genuine neural signals. Therefore, we assessed the robustness of our results using both uncorrected data and data corrected with a fourth-order polynomial detrending approach (13), in order to account for potential preprocessing-related distortions. Functional connectivity was estimated for each condition by computing pairwise Pearson correlation coefficients between oxy-Hb time series from all channel combinations. Prior to this, a band-pass filter (0.009–0.1 Hz) was applied to isolate spontaneous low-frequency oscillations associated with functional connectivity.

### Statistical Analysis

A linear mixed-effects model was used to compare swimming performance (total 100 m time) between conditions. The model included a fixed effect of condition (tDCS vs. sham) and a random intercept for each participant to account for repeated measures. For the fNIRS data, we used linear mixed-effects models to analyze changes in both cortical activation (oxy-Hb) for each of the 14 channels and functional connectivity for each of the 91 channel pairs. These models included fixed effects for time (baseline vs. stimulation), condition (tDCS vs. sham), and their interaction term (time × condition). A random intercept for each participant was included. To optimize model convergence, we employed a stepwise approach for selecting the random effects structure. We began with a maximal model including random slopes for all fixed effects and progressively simplified it if the model failed to converge or resulted in a singular fit. P-values for the multiple fNIRS analyses (14 for activation, 91 for connectivity) were corrected using the false discovery rate (FDR) procedure. To obtain overall patterns of the results, we additionally grouped channel pairs into intra- and inter-hemispheric categories—left-left (L-L), left-right (L-R), and right-right (R-R)—based on their anatomical locations. For these groups, we computed the average standardized regression coefficients. Although post-hoc tests were performed only when significant interactions were detected at the channel-pair level, effect sizes for all three categories were calculated to provide a comprehensive overview of the connectivity patterns. Finally, Pearson’s correlation analyses were used to examine the relationship between neural measures (functional connectivity) and behavioral outcomes (swimming time), both within the tDCS condition and for the difference scores (tDCS versus sham). Given the exploratory nature of these analyses with a small sample, correlation coefficients are reported with their 95% confidence intervals (CIs). All analyses were performed in R (version 4.5.0).

## RESULTS

### Participant Characteristics

On average, participants were 20.5 ± 1.2 years old and had 15.1 ± 6.3 years of competitive swimming experience. They trained approximately 3.5 ± 2.7 days per week, with a daily training duration of 1.5 ± 0.9 hours. The mean height and weight were 171.0 ± 7.1 cm and 61.3 ± 4.5 kg, respectively. The participants’ personal best times in the 100 m freestyle (short-course; 25 m pool) averaged 58.57 ± 10.64 seconds.

### Manipulation Check

The blinding procedure was successful. There were no significant differences between conditions in terms of sleepiness (tDCS vs. sham: 5±1 vs. 4±1, Cohen’s d = 0.41, p = 0.28), itching (tDCS vs. sham: 50% vs. 63%, χ^2^ = 0.00, p = 1.00), pain (25% vs. 38%, χ^2^ = 0.00, p = 1.00), burning sensation (13% vs. 0%, χ^2^ = 0.00, p = 1.00), warmth (38% vs. 38%, χ^2^ = 0.00, p = 1.00), metallic/iron taste (none reported), fatigue (38% vs. 38%, χ^2^ = 0.00, p = 1.00), and other symptoms (13% vs. 0%, χ^2^ = 0.00, p = 1.00). Additionally, there were no significant differences in the timing of sensations (beginning: 38% vs. 38%, χ^2^ = 0.00, p = 1.00; middle: 63% vs. 50%, χ^2^ = 0.00, p = 1.00; end: 0% vs. 25%, χ^2^ = 0.57, p = 0.45), their duration (beginning only: 25% vs. 38%, χ^2^ = 0.00, p = 1.00; up to the middle: 88% vs. 50%, χ^2^ = 1.16, p = 0.28; to the end: 25% vs. 25%, χ^2^ = 0.00, p = 1.00), subjective assessments of impact on overall condition (1.88 vs. 1.63, t_7_ = 1.00, p = 0.35), or location (throughout: 38% vs. 13%, χ^2^ = 0.33, p = 0.56; partially: 38% vs. 63%, χ^2^ = 0.25, p = 0.62; near electrode: 13% vs. 13%, χ^2^ = 0.00, p = 1.00) during stimulation.

### Physical Performance

The total 100 m freestyle time was numerically shorter in the tDCS condition compared to the sham condition (62.8 s vs. 63.2 s), but this difference was not statistically significant (mean difference [95% CI] = −0.44 s [−1.54, 0.66], p = 0.38) (Figure 2A). An exploratory analysis of the four 25-meter segments revealed that swim times were numerically shorter in the tDCS condition for all segments. This difference was most pronounced in the second segment (25–50 m), where performance under tDCS was faster than under sham (15.8 s vs. 16.0 s; mean difference [95% CI] = −0.28 s [−0.55, −0.01]). This segment-specific improvement was statistically significant at an uncorrected level (p = 0.04), though it did not survive correction for multiple comparisons (Figure 2B).

**Figure 2.**
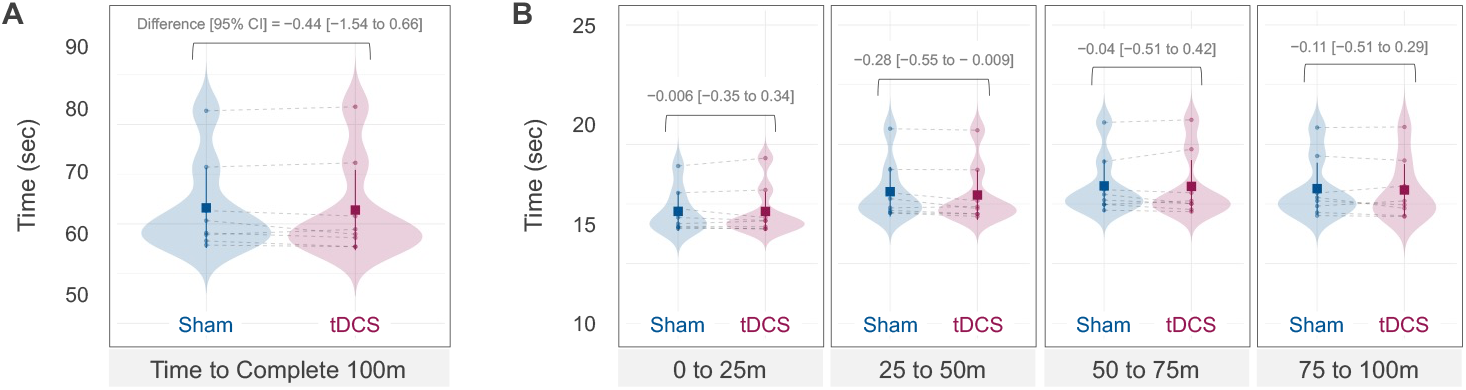
Swimming performance under tDCS and sham conditions. A. Total time (in seconds) to complete the 100-meter freestyle swimming task. Violin plots show the distribution of times under sham (blue) and tDCS (red) conditions, with mean values (points) and 95% confidence intervals (error bars). B. Completion times are broken down by 25-meter segments (0–25 m, 25–50 m, 50–75 m, and 75–100 m). Each panel compares tDCS and sham conditions, with corresponding violin plots and 95% CIs. The estimated difference and 95% CI from linear mixed-effects models are indicated at the top of each panel.

### Hemodynamic Response

The time courses of hemodynamic changes and functional connectivity matrices are shown in Figure 3. Although oxy-Hb levels showed a slight increase during stimulation in both conditions, this trend was attenuated following baseline drift correction. No significant condition × time interaction effects on cortical activation were observed for any channel after applying FDR correction (ps > 0.30). In contrast, the analysis of functional connectivity revealed significant effects. As shown in the heatmaps (Figure 4A), we observed significant condition × time interactions, primarily reflecting a decrease in connectivity during stimulation in the tDCS condition. These effects were concentrated in intra-hemispheric connections within the stimulated left hemisphere (Figure 4B). Significant interactions were confirmed for the connectivity between channels 7–8 (β = −1.40, FDR-adjusted p = 0.02) and channels 10–11 (β = −1.51, FDR-adjusted p = 0.02). Post-hoc analyses of these significant interactions (Figure 4C) revealed that in the tDCS condition, functional connectivity significantly decreased from baseline to stimulation for both channel pairs (7–8: β = −1.00, FDR-adjusted p < 0.001; 10–11: β = −0.72, FDR-adjusted p = 0.006). Conversely, in the sham condition, connectivity significantly increased for both pairs (7–8: β = 0.40, FDR-adjusted p = 0.04; 10– 11: β = 0.79, FDR-adjusted p = 0.003). Direct comparison during the stimulation period showed significantly lower connectivity in the tDCS condition compared to the sham condition for channels 7–8 (β = −1.23, FDR-adjusted p = 0.02). However, no significant difference was found between the conditions for channels 10-11 (β = −0.73, FDR-adjusted p = 0.07).

**Figure 3.**
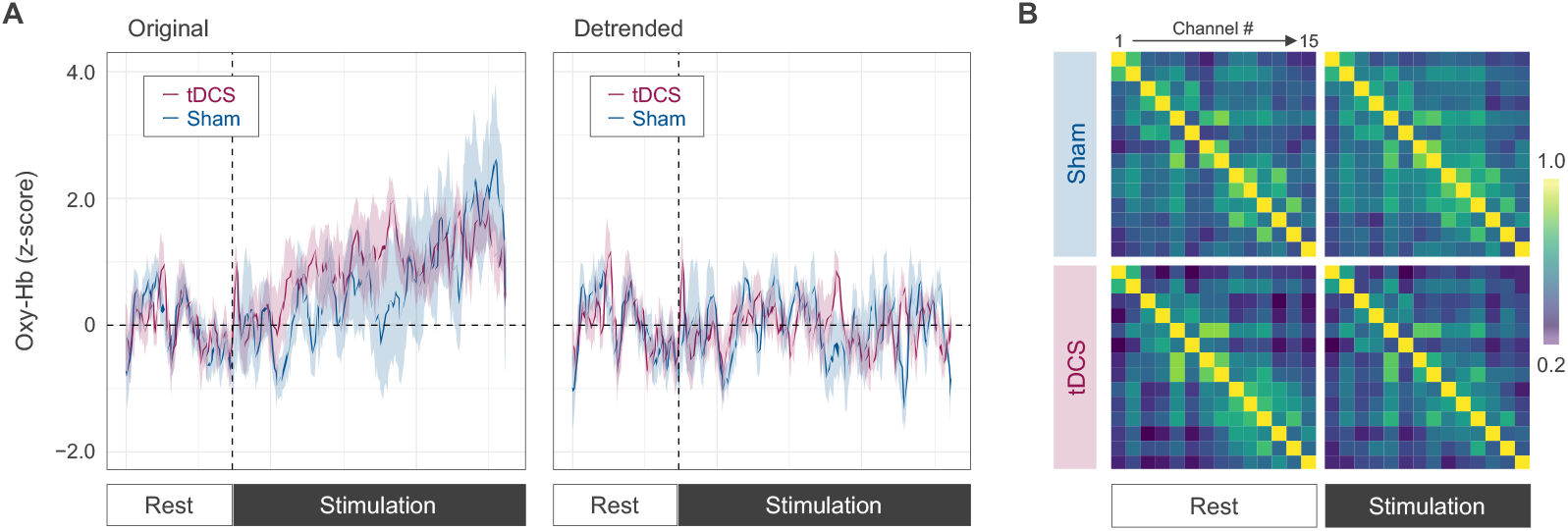
Hemodynamic response and functional connectivity during rest and stimulation phases. A. Time series of z-scored changes in oxygenated hemoglobin (Oxy-Hb) averaged across all fNIRS channels. Left panel shows the original data; right panel shows the data after detrending. Solid lines represent the mean for each condition (red: tDCS, blue: sham), and shaded areas indicate 95% confidence intervals. The vertical dashed line marks the onset of the stimulation phase. B. Group-averaged functional connectivity matrices derived from Pearson’s correlation coefficients between channel pairs, shown separately for rest and stimulation phases under sham (top) and tDCS (bottom) conditions.

**Figure 4.**
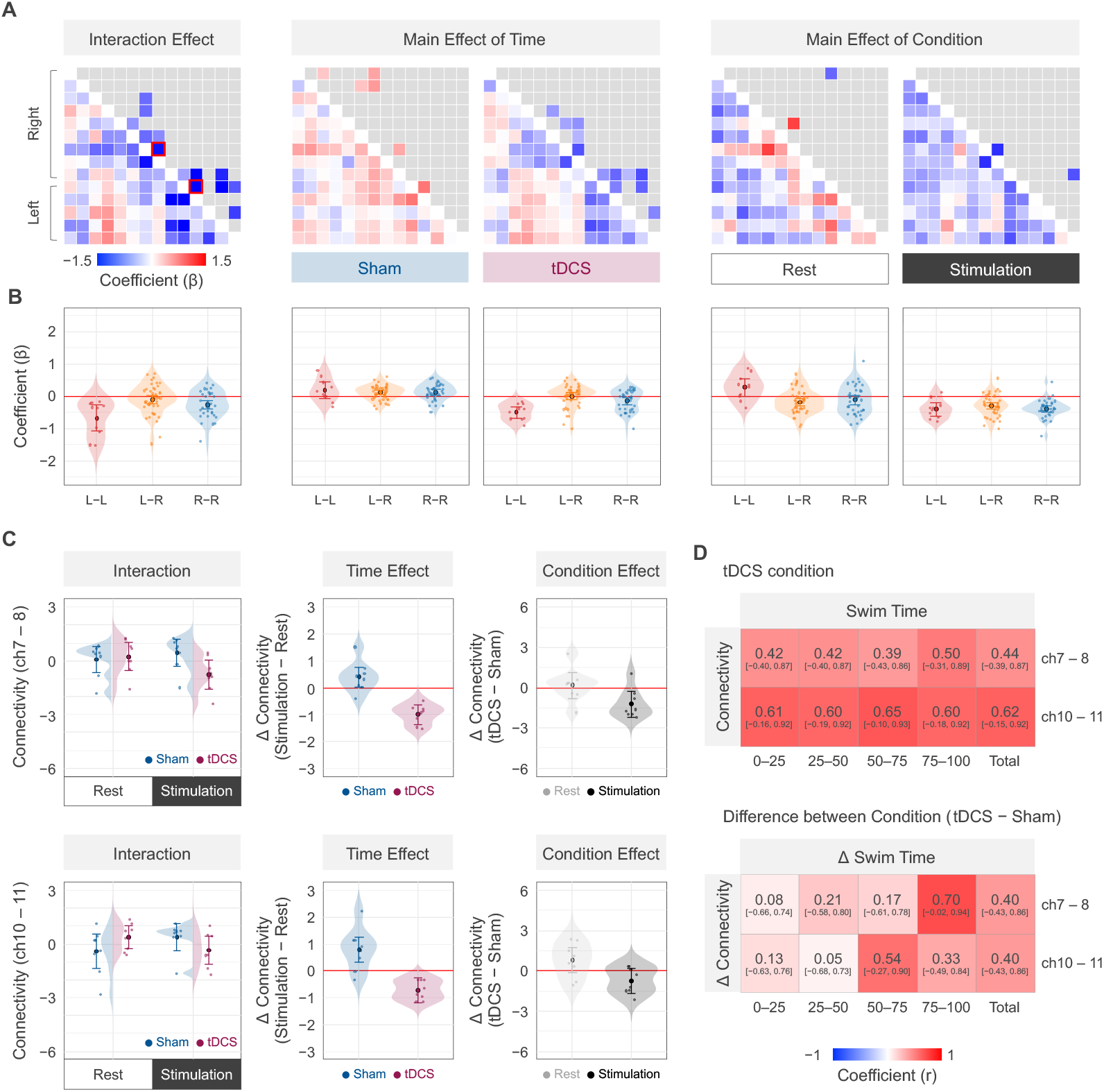
Linear mixed-effects model and correlation results. A. Heatmaps of standardized regression coefficients (B values) showing the interaction between condition and time (left), main effects of time within each condition (middle), and main effects of condition at each time point (right). Note that for visualization purposes, channel pairs showing uncorrected p-values < 0.05 are marked in the upper triangle of the heatmaps, but no statistical inference should be drawn from these markings. Red outlines highlight regions with significant interaction effects after correction for multiple comparisons (channel pairs 7–8 and 10–11). B. Violin plots of B estimates for intra-hemispheric (L-L, R-R) and inter-hemispheric (L-R) connectivity for each effect. Midline channels (i.e., channel pair 8–9) are included in the right. C. Plots of interaction effects and post-hoc analysis in specific channel pairs (7–8 and 10–11) that showed significant condition-by-time interactions after FDR correction. Error bars denote 95% confidence intervals. D. Heatmaps of Pearson’s correlation coefficients between functional connectivity and performance. The upper panel shows within-condition (tDCS only) correlations, while the lower panel shows correlations between the differential effects of tDCS (tDCS – Sham) on connectivity and performance.

### Relationship Between Neural and Behavioral Outcomes

We next examined the relationship between neural and behavioral outcomes. Within the tDCS condition, there was a positive correlation between the functional connectivity during stimulation and the 100 m swim time. This indicates that lower functional connectivity was associated with better performance (i.e., a shorter swim time). Correlation coefficients for channel pairs within the left PFC ranged from r = 0.39 to r = 0.65 (Figure 4D).

Furthermore, we examined the relationship between difference scores (tDCS–sham) for both connectivity and performance. This analysis also revealed a positive correlation between the difference scores for connectivity and performance, with coefficients ranging from 0.09 to 0.70 (Figure 4D). Specifically, a greater reduction in functional connectivity under tDCS (relative to sham) was associated with a greater improvement in swimming time (relative to sham).

### Exploratory Analysis

To further investigate the potential source of individual differences in responsiveness to tDCS, we conducted an exploratory analysis. Based on visual inspection of the data, we hypothesized that baseline levels of functional connectivity during rest could predict the degree of connectivity change induced by tDCS. This was suggested by the observation that channel pairs with stronger functional connectivity at rest in the tDCS condition tended to show greater reduction in connectivity during tDCS (Figure 4A). We formally tested this by correlating the main effect of condition at rest with the main effect of time in the tDCS condition, revealing a strong negative correlation (r = −0.50, 95% CI [−0.63, −0.34]).

Based on this, we conducted a follow-up exploratory analysis at the individual participant level. We examined the correlation between each individual’s functional connectivity at rest and their change in connectivity from rest to stimulation, separately for each condition. The results showed a strong negative correlation in the tDCS condition (mean r = –0.57, 95% CI [–0.61, –0.53]), indicating that participants with higher baseline connectivity experienced larger decreases during stimulation. In contrast, the sham condition showed only a weak negative correlation (mean r = –0.25, 95% CI [–0.29, –0.21]).

To distinguish between stable between-subject differences (trait) and transient within-subject fluctuations (state), we further examined the correlation between resting-state functional connectivity measured in the tDCS and sham sessions. This analysis yielded a moderate positive correlation (r = 0.43, 95% CI [0.39, 0.46]), suggesting that individual differences in baseline functional connectivity are at least partially stable across sessions. This supports the interpretation that baseline connectivity may reflect a trait-like characteristic that moderates responsiveness to tDCS. However, the moderate magnitude of the correlation also indicates that session-to-session variability may contribute to tDCS effects, implying the involvement of both trait and state components.

## DISCUSSION

This study investigated the neurobehavioral effects of prefrontal tDCS on swimming performance by combining fNIRS neuroimaging with a 100 m time-trial task. Our results offer several key insights. First, while tDCS over left DLPFC produced only a small, non-significant overall improvement in 100 m freestyle time, it appeared to enhance performance in the specific 25–50 m segment. Second, at the neural level, tDCS did not alter overall prefrontal activation but significantly decreased functional connectivity, especially within the stimulated left hemisphere. Most importantly, these two findings were linked: the magnitude of the connectivity decrease correlated with the degree of performance enhancement. Finally, our exploratory and follow-up analyses suggest that both stable between-subject (trait-like) and transient within-subject (state-like) factors may contribute to individual responsiveness to tDCS.

Regarding performance, the lack of a statistically significant improvement in the overall 100 m time is consistent with the broader tDCS literature, which is characterized by inconsistent results and small effect sizes, likely due to high inter-individual variability (5). This lack of a significant group-level effect is consistent with the study’s premise that substantial inter-individual variability is a key factor influencing outcomes (6–11). Given the substantial inter-individual variability in neural responses (6–11), a significant average improvement is unlikely unless most participants are responders. The overall null result, therefore, likely reflects the performance gains in a subset of individuals being diluted by the lack of change in others. Nevertheless, the finding that tDCS improved performance in the 25–50 m segment is intriguing. This segment follows the initial explosive start and turn, and precedes the onset of major fatigue. The PFC is crucial for cognitive control essential for elite performance, such as executing a pacing strategy, maintaining attentional control, and managing the perception of effort (28). It is plausible that tDCS did not enhance maximal power output but instead optimized mid-race pacing or reduced the cognitive load of maintaining technique, thereby enabling a segment-specific gain. This aligns with previous studies in swimmers that found positive effects of tDCS on shorter sprints (e.g., 50 m) but null effects on longer or all-out swims (20–22). While this finding is potentially hypothesis-generating, it should be interpreted with great caution as it is not statistically robust and did not withstand correction for multiple comparisons.

Second, our primary neuroscientific finding was that tDCS modulated functional connectivity rather than overall cortical activation. Specifically, we observed a significant decrease in intra-hemispheric connectivity within the stimulated left PFC. This contributes to the ongoing debate on tDCS’s effects on network properties. A decrease in connectivity, particularly in experts, is often interpreted as a marker of neural efficiency (29). In this framework, an efficient neural network requires less functional coupling between regions to perform a task. While anodal tDCS is generally considered excitatory at the cellular level, its network-level effect may be to optimize neural processing by reducing redundant co-activation and enhancing computational efficiency. This enhancement of neural efficiency in the PFC could be a key physiological mechanism for mitigating the rise in perceived exertion and delaying central fatigue, thereby allowing athletes to better sustain motor output and technical precision during demanding tasks.

The core finding of this study is the direct brain-behavior link we established: athletes whose prefrontal networks showed a greater decrease in connectivity also swam faster. This finding suggests a crucial corollary: the inconsistency in the tDCS literature may stem from a failure to account for variability in neural responses. Our data imply that individuals can be categorized as neural responders and non-responders, and that failing to measure the underlying neural effects may lead to diluted group-level behavioral results. By integrating neuroimaging, our study provides a potential mechanistic explanation for the mixed results that have long plagued tDCS research (6, 7). Building on this brain-behavior link reinforces the neural efficiency hypothesis (29). The top-down control exerted by the PFC is vital in elite sports (28), and enhancing its efficiency appears to be a key mechanism of tDCS.

Our exploratory analysis further suggests a “normalization” effect: individuals with higher baseline connectivity—potentially indicating a less efficient or more noisy network at rest—benefited most from the stimulation. This aligns with recent proposals that tDCS can “normalize” inefficient brain activation and cortical networks (30, 31). tDCS may act to prune redundant connections, sharpening network function. This effect is naturally most pronounced in those who have the most room for improvement in neural efficiency. While the weak negative correlation in the sham condition could reflect simple regression to the mean, the much stronger correlation in the tDCS condition points to a genuine physiological effect, not a mere statistical artifact.

To further investigate whether tDCS effects were primarily driven by between-subject or within-subject variability in baseline connectivity, as an exploratory step, we evaluated the correlation between resting-state connectivity values across the two sessions. The moderate correlation (r = 0.43) suggests that resting-state connectivity has both stable and variable components. This indicates that while some individual differences in responsiveness to tDCS reflect stable neural architecture, transient factors such as fatigue, motivation, or recent training load may also modulate tDCS responsiveness. This has profound practical implications. It suggests that a one-size-fits-all approach to tDCS is suboptimal. Future protocols could involve baseline neuroimaging to screen for likely “responders,” allowing for a personalized approach to maximize the intervention’s reliability as a performance-enhancement tool. For example, one might envision athletes undergoing pre-competition fNIRS scans to tailor neuromodulation to their specific neural state on a given day.

Several limitations of this study must be acknowledged. The primary limitation is the small sample size (n = 8), which limits the generalizability of our findings and increases the risk of Type II errors.

Therefore, our results should be considered preliminary and as a proof-of-concept. However, it is important to note that the rigorous double-blind, sham-controlled, crossover design significantly enhanced internal validity by allowing each participant to serve as their own control. This design is particularly powerful for studying interventions like tDCS, where inter-individual variability is a major confounding factor. Our focus on individual-level predictors, rather than solely on group-level averages, represents a crucial step toward understanding this variability. Second, fNIRS, while ecologically valid, is an indirect measure of neural activity and has lower spatial resolution and penetration depth than fMRI. Finally, we investigated the acute effects of a single tDCS session. Future research should build upon these findings by recruiting larger, more diverse cohorts (including female athletes and athletes from different sports) and by exploring the cumulative effects of multiple sessions, which may be required to induce lasting neuroplasticity and more physiologically meaningful and statistically robust performance gains.

In conclusion, this study provides novel evidence that anodal tDCS over the left DLPFC modulates functional connectivity in competitive swimmers. Although a significant improvement in overall 100 m performance was not detected, the degree of connectivity reduction correlated with performance gains. This suggests that tDCS may enhance physical performance by increasing the neural efficiency of prefrontal networks. Moreover, our findings indicate that baseline connectivity may predict an individual’s response to stimulation, paving the way for a more personalized, biomarker-guided approach to neuromodulation in high-performance settings. While the current work is exploratory, it offers a foundational step toward unlocking the potential of tDCS by providing a mechanistic framework and identifying a tangible biomarker to guide its application in sports.

## ACKNOWLEDGMENTS

We thank Masanobu Homma, Yoshiharu Inou, and Kohei Masumoto for their help in conducting this study.

## Notes

### Competing Interest Statement

The authors have declared no competing interest.

